# Cassava witches’ broom disease in French Guiana: a threat to cacao cultivation and its biodiversity?

**DOI:** 10.64898/2026.04.05.716555

**Authors:** Abdoul-Raouf Sayadi Maazou, Fabien Doare, Eliane Louisanna, Hélène Vignes, Didier Tharreau, Henri Adreit, Clara Cayron, G. Martijn Ten Hoopen

## Abstract

Beyond the significant impact of Cassava witches’ broom disease (CWBD), caused by the fungus *Rhizoctonia (*syn. *Ceratobasidium) theobromae* on cassava cultivation in French Guiana and Brazil, this disease also poses a potential threat to cacao trees in the region, since the fungus is responsible for Vascular Streak Dieback (VSD) of cacao in South East Asia. Cross-pathogenicity trials were conducted in several cassava fields in French Guiana by planting young cacao plants adjacent to diseased cassava plants. Vascular necrosis was observed in some cacao plants, and the presence of *R. theobromae* in the cacao tissues was confirmed through PCR diagnostics using primers specific to the fungus. Sequence analysis indicated 100% similarity between samples from both hosts and 97.53 to 99.74% identity with *R. theobromae* isolates previously reported from cassava in the Americas and Southeast Asia. Additionally, symptomatic cacao in a mixed cacao–cassava farm yielded *R. theobromae*–positive PCR results, suggesting a natural infection. Ongoing work includes artificial inoculations and controlled cross–pathogenicity trials under screenhouse conditions to attempt reproduction of the symptoms. While current data do not yet establish definitive causality, the findings indicate potential host jump and warrant rapid communication to researchers, policy makers, and farmers to safeguard cacao production and *Theobroma* biodiversity in the Amazon region.

---

Cassava witches’ broom disease (CWBD) caused by the fungus *Rhizoctonia theobromae* (syn. *Ceratobasidium theobromae*) has long been recognized as a major transboundary disease in Southeast Asia (Pardo et al., 2023), posing a significant challenge for cassava (*Manihot esculenta* Crantz) production in the region. After several years of confinement in southeast Asia, the disease has been reported in the Americas (Pardo et al., 2024). It was first detected in French Guiana around 2021/2022, near Maripasoula, before spreading to almost all municipalities of the territory. Since cassava is a major source of food for many local communities, this disease seriously threatens their food security. It alters their eating habits, reduces their income, and exacerbates the already precarious situation for many Indigenous communities. A similar pattern has also been reported in Brazil (Oliveira et al., 2025).

In addition to the devastating impact of CWBD on cassava cultivation in French Guiana and Brazil, *Rhizoctonia theobromae* also poses a potential threat to cacao (*Theobroma cacao* L.) trees in the region, since *R. theobromae* is responsible for vascular Streak Dieback of cacao in South East Asia (McMahon & Purwantara, 2016). Both diseases share common symptoms such as vascular tissue necrosis, leaf drop, and the development of axillary buds, resulting in the formation of small, broom-like branches in cacao trees. *Rhizoctonia theobromae* spores, generally wind-borne, infect young, soft leaves at the branch tips and colonize the leaf xylem. From the leaf, the fungus reaches the stem via the petiole and spreads through the xylem vessels, causing vascular necrosis and branch dieback, sometimes killing young cacao trees (Marelli et al., 2019). Phylogenetic analyses revealed that isolates of the pathogen causing CWBD are very closely related but genetically distinct of isolates causing vascular streak dieback (VSD) in cacao (Leiva et al., 2023; Tobias et al., 2026). Field observations suggest a similar mode of transmission for CWBD and VSD, which could indicate that *R. theobromae* could infect both cassava and cacao. The disease could therefore spread from contaminated cassava fields to nearby cacao plantations. Moreover, the question of whether *R. theobromae* infecting cacao could also infect other species of the genus *Theobroma* or the closely related genus *Herrania* remains unresolved. Considering such risks, we initiated cross-pathogenicity tests in French Guiana, intending to verify the ability of the fungus *R. theobromae*, present on cassava plants, to infect cacao plants and produce the VSD symptoms.

Three cassava fields in Iracoubo municipality, French Guiana, each infected with CWBD and located well away from any cacao plantation, were selected for a cross-pathogenicity trial. GPS coordinates for the three fields were 5°28′45.097″N, 53°12′14.21″W; 5°28′49.117″N, 53°12′16.024″W; and 5°28′45.123″N, 53°12′13.934″W. We obtained 120 cacao plants (two groups of 60), representing open-pollinated offspring of Pound 7 and IMC 97 clones, from the Biological Resource Centre for Perennial Plants in French Guiana (CRB-PPG). In March 2025, these plants were transplanted adjacent to diseased cassava plants in the three fields. Planting density in each field was 1 m × 1 m, arranged in three blocks per field; each block contained 10 cacao plants (five Pound 7 and five IMC 97). An additional control block was established ∼3 km from the infected fields.

Cacao growth was monitored for five months, after which samples were collected for PCR diagnostics following observation of necrotic stem tissues on eight cacao plants. From each plant, a 5–8 cm apical stem cut (affected tissue in symptomatic plants) was collected, split longitudinally, and vascular tissue scraped for DNA extraction using a CTAB protocol (Doyle, 1991). PCR targeting the Ca2+/calmodulin-dependent protein kinase gene (CaMK) of *R. theobromae* (GenBank accession number KAB5596398) with the primer reported by Leiva et al. (2023) amplified the target sequence in eight cacao plants exhibiting vascular necrosis (Fig. 1) and in 16 cassava samples collected from the same fields. The positive cacao samples were collected from two of the three fields. No symptoms were observed on cacao plants of the control blocks, and control samples were PCR-negative. The PCR diagnostics are summarized in Table 1, and agarose gel electrophoresis images are presented in Supplementary file 1 (ESM 1). Positive PCR products from cacao and cassava samples were Sanger sequenced (GenoScreen), and the resulting sequences were analyzed using BLASTn. A sequence from each host and field was deposited in GenBank and provided in Supplementary file 2 (ESM 2). Cacao samples are CAMK GUF□C7 (GenBank accession no. PZ112601) and CAMK GUF□C98 (PZ112602); cassava samples are CAMK GUF□IA37 (PZ112598), CAMK GUF□IB17 (PZ112599), and CAMK GUF□IC10 (PZ112600). The cacao and cassava sequences were 100% identical (excluding missing nucleotides) and showed 97.53–99.74% identity to *R. theobromae* isolates previously reported from cassava in French Guiana, Brazil, Laos, and Vietnam (GenBank accession nos. PQ384454, PQ202694, OQ863083, OQ863064).

**Table 1.** PCR analysis of cassava and cacao stem samples collected from cross□pathogenicity trial implemented in three cassava fields in Iracoubo municipality, French Guiana.

| Sample type | Number of samples | PCR (-) | PCR (+) |
| --- | --- | --- | --- |
| Healthy cassava | 28 | 27 | 1 |
| Diseased cassava | 92 | 68 | 24 |
| Healthy cacao | 112 | 112 | 0 |
| Diseased cacao | 8 | 0 | 8 |

**Fig. 1.**
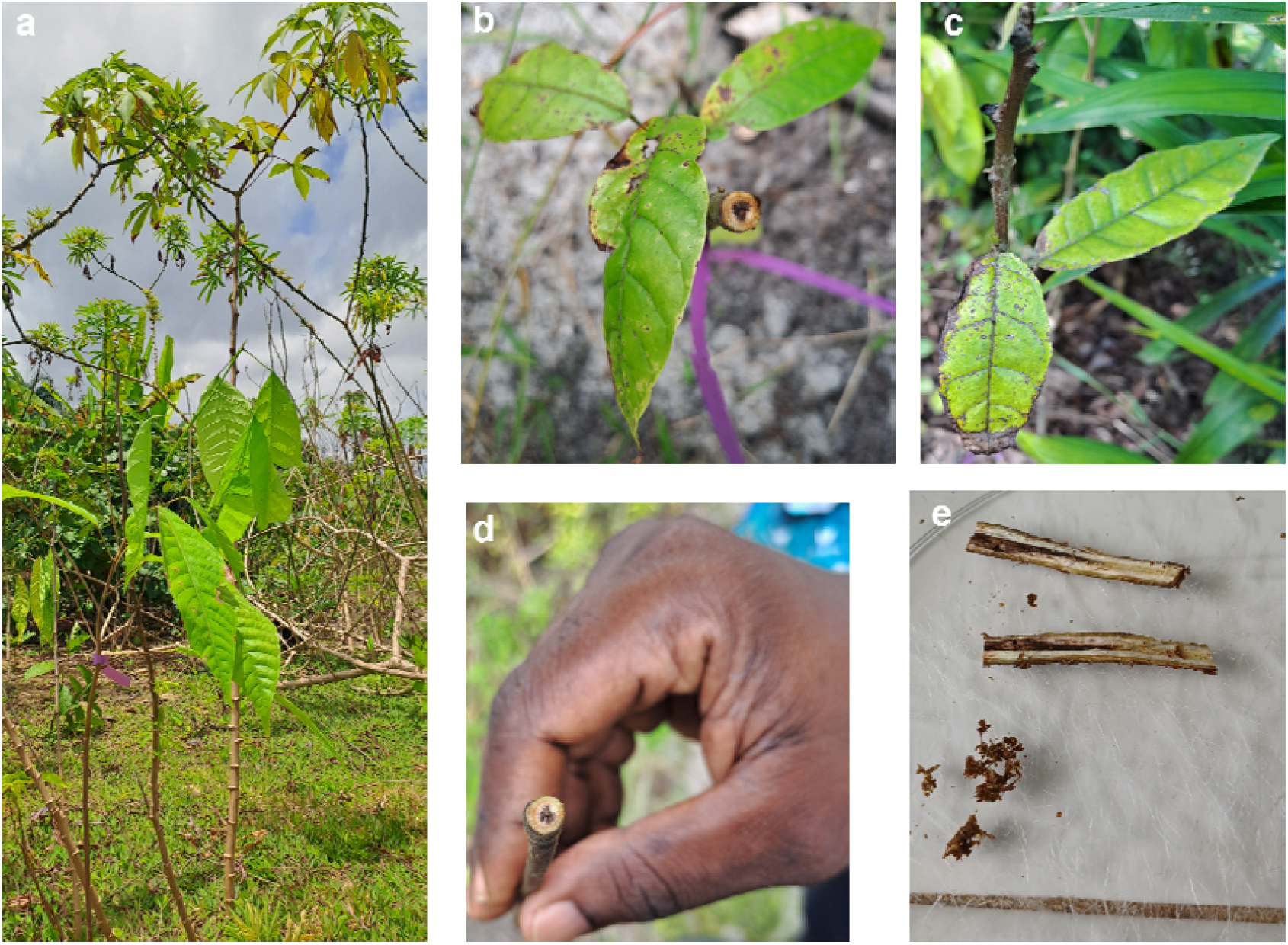
Symptoms of *Rhizoctonia theobromae* infection on *Theobroma cacao* L. from a cassava–cacao cross□pathogenicity trial in French Guiana. **a**. Field view of one of the cross□pathogenicity plots showing cacao plants grown at high density beneath cassava plants exhibiting cassava witches’ broom symptoms; **b**. Cross section of a cacao stem showing vascular necrosis; **c**. Leaf with necrotic lesions; **d**. Stem sample collected for PCR diagnosis; **e**. Longitudinal section of an infected stem with scraped central vascular tissue used for molecular analysis

In addition to the cross-pathogenicity trials, during routine surveillance, FREDON Guyane reported CWBD-like symptoms on cassava in a mixed cacao–cassava farm in Roura municipality, French Guiana (4°41′23.712″N, 52°24′9.54″W) in December 2025. A follow-up visit within days confirmed extensive necrosis on cacao (Fig. 2) and typical CWBD symptoms on cassava, including chlorosis, shortened internodes, vascular necrosis, and brooming (weak, spindly sprouts) on stems. Samples for DNA extraction and PCR were collected from both hosts; extraction and amplification followed the procedures described above. Of 10 cacao and 5 cassava samples, the *R. theobromae* CAMK sequence consistently amplified in 1 cacao and 4 cassava samples, suggesting a natural infection of cacao plants (Fig.3). This is the first report of R. theobromae infecting cacao in French Guiana.

**Fig. 2.**
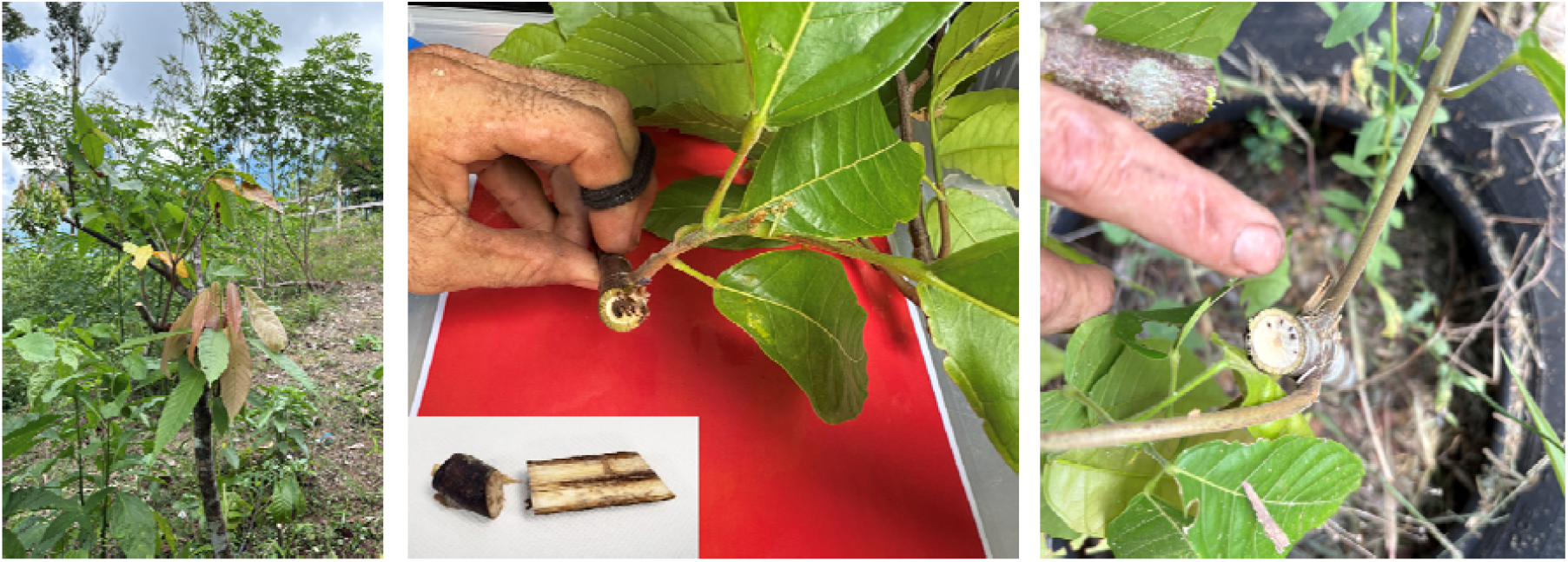
Symptoms of *Rhizoctonia theobromae* infection on *Theobroma cacao* L. under natural infection conditions in a mixed cacao–cassava farm in French Guiana

**Fig. 3.**
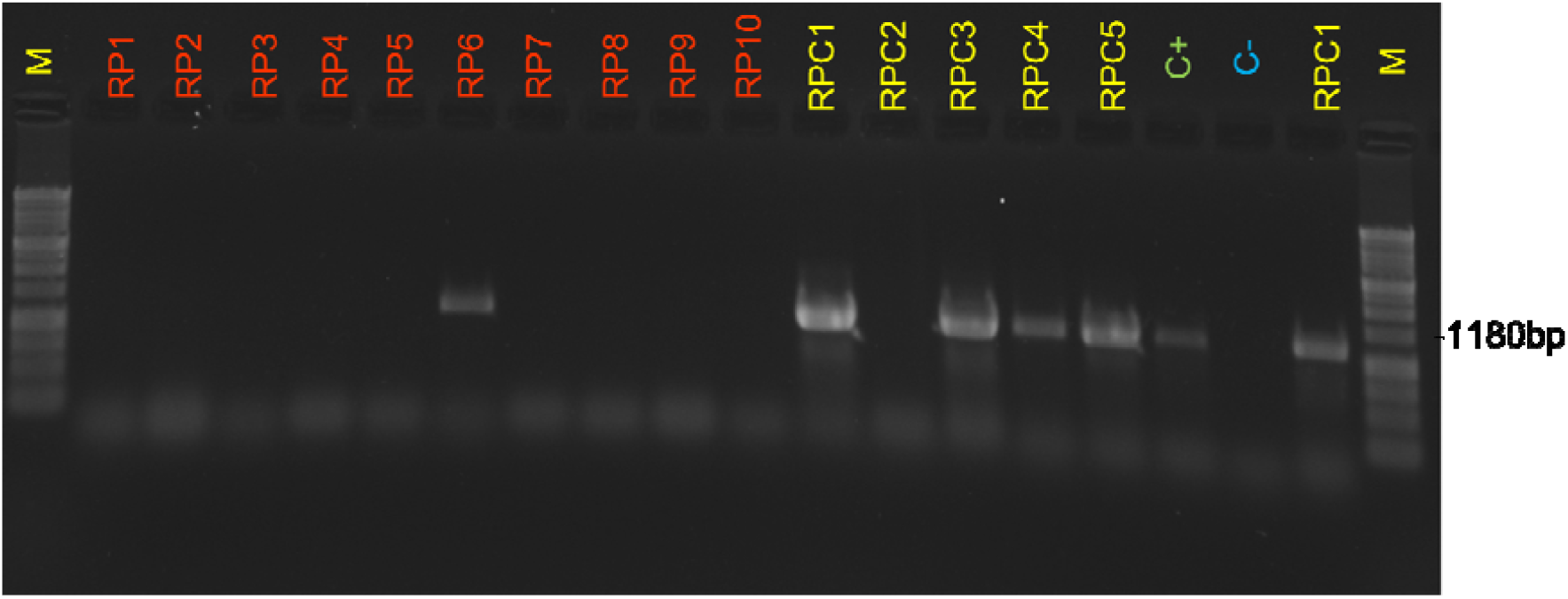
Agarose gel electrophoresis showing Rhizoctonia theobromae PCR amplification (target size 1180bp). **Lane M**: 1kb DNA ladder**; Lanes RP1-RP10**: cacao samples**; Lanes RPC1-RPC5**: cassava samples; **Lane C+**: positive control (DNA extracted from a symptomatic cassava plant); **Lane C-**: negative control

While these results indicate a potential host jump with implications for cacao, they do not yet satisfy Koch’s postulates. It therefore remains uncertain whether *R. theobromae* strains affecting cassava can cause disease in *Theobroma* spp. Ongoing work includes artificial inoculation of young cacao plants with mycelium from cassava isolates and controlled cross□pathogenicity trials in screenhouse conditions to determine whether the fungus can infect cacao and reproduce the symptoms of VSD. Meanwhile, it is urgent to promptly inform the scientific community and relevant policy makers and raise awareness among farmers. Until pathogen transmission is clarified, planting cassava adjacent to cacao should be avoided, and intensified monitoring in French Guiana and Brazil should be pursued to enable early detection and guide development of management and prevention strategies to protect cacao production and *Theobroma* biodiversity in the Amazon region.

## Acknowledgements

We thank the field technician Miguel Adrian of the ‘‘Association des Agriculteurs des Savanes (ADADS)’’ for his support in the implementation of the field trials, field visits, and sample collection. We also express our gratitude to Valérie Troispoux of the UMR EcoFoG Genetic laboratory for help during sample processing, and to the cassava farmers of Iracoubo municipality, French Guiana, who kindly allowed us to visit their fields and implement the trials.

## Funding

This work has been funded by the French Government through the “Direction générale de l’alimentation (DGAL)” of ‘‘Ministère de l’Agriculture, de l’Agro-alimentaire et de la Souveraineté alimentaire’’ under the projects DECODE (C-2024-121) and DECODE+ (CIRAD-01).

## Authors contribution

Conceptualization: Ten Hoopen GM, Maazou A-RS, Tharreau D. Investigation: Maazou A-RS, Doare F, Louissanna E, Vignes H, Tharreau D, Cayron C, Adreit H. Writing: Maazou A-RS. Review and Editing: Doare F, Vignes H, Tharreau D, Cayron C, Adreit H, Ten Hoopen GM. Project administration: Maazou A-RS. Funding Acquisition: Ten Hoopen GM, Maazou A-RS.

## Data availability

All data supporting the findings of this study are available within the paper and its supplementary files.

## Declarations

### Competing interest

The authors declare no competing interests.

## Notes

### Competing Interest Statement

The authors have declared no competing interest.

