## Supplementary file 1 for "Cassava witches’ broom disease in French Guiana: a threat to cacao cultivation and its biodiversity?"

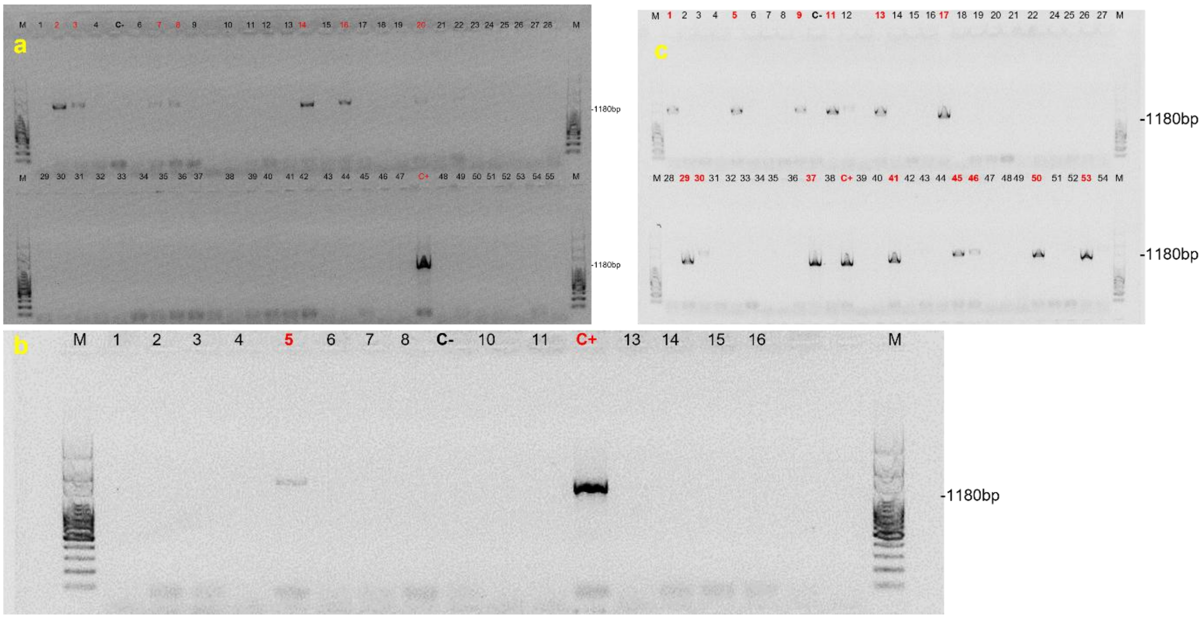


Supplementary Figure S1. Agarose gel electrophoresis showing *Rhizoctonia theobromae* PCR amplification (target size 1180bp). **a. Lane M**: 1kb DNA ladder**; Lanes 1-55**: cacao samples (GUF C1 – C55); **Lane C-:** negative control; **Lane C+**: positive control (DNA extracted from a symptomatic cassava plant); **b. Lane M**: 1kb DNA ladder**; Lanes 1-16**: cacao samples (GUF C94 – C107); **Lane C-:** negative control; **Lane C+**: positive control; **c. Lane M**: 1kb DNA ladder**; Lanes 1-54**: cassava samples (GUF IA1 – IB14); **Lane C-:** negative control; **Lane C+**: positive control
